# Repurposing Hsp90 inhibitors as antimicrobials targeting two-component systems identifies compounds leading to loss of bacterial membrane integrity

**DOI:** 10.1101/2023.12.13.571547

**Authors:** Blanca Fernandez-Ciruelos, Marco Albanese, Anmol Adhav, Vitalii Solomin, Arabela Ritchie-Martinez, Femke Taverne, Nadya Velikova, Aigars Jirgensons, Alberto Marina, Paul W. Finn, Jerry M. Wells

## Abstract

The discovery of antimicrobials with novel mechanisms of action is crucial to tackle the foreseen global health crisis due to antimicrobial resistance. Bacterial two-component signalling systems (TCS) are attractive targets for the discovery of novel antibacterial agents. TCS-encoding genes are found in all bacterial genomes and typically consist of a sensor histidine kinase (HK) and a response regulator (RR). Due to the conserved Bergerat fold in the ATP-binding domain of the TCS HK and the human chaperone Hsp90, there has been much interest in repurposing inhibitors of Hsp90 as antibacterial compounds. In this study, we explore the chemical space of the known Hsp90 inhibitor scaffold 3,4-diphenylpyrazole (DPP), building on previous literature to further understand their potential for HK inhibition. Six DPP analogues inhibited HK autophosphorylation *in vitro* and had good antimicrobial activity against Gram-positive bacteria. However, mechanistic studies showed that their antimicrobial activity was related to damage of bacterial membranes. In addition, DPP analogues were cytotoxic to mammalian cancer cell lines and induced the cell arrest phenotype shown for other Hsp90 inhibitors. We conclude that these DPP structures can be further optimized as specific disruptors of bacterial membranes providing binding to Hsp90 and cytotoxicity are lowered. With respect to the original hypothesis, the X-ray crystal structure of resorcinol, a substructure of the DPP derivatives, bound to the HK CheA represents a promising starting point for the fragment-based design of novel HK inhibitors.

## Introduction

The rise of antibiotic resistance worldwide makes it necessary to find new antimicrobial treatments, preferably with low potential for resistance development(1). Two-component systems (TCSs), the most important signalling systems in bacteria, are promising antibacterial targets(2). They are absent in animal cells(3) and present in all bacteria(4). Each bacterial species encodes multiple TCSs(5–7) involved in adaptive responses and the regulation of metabolism(8), response to extracellular stresses(9), antibacterial resistance(10), and virulence in the host(11), among others. Typically, TCSs consist of a sensor histidine kinase (HK) and a response regulator (RR)(12). Prototypical HKs are membrane-bound dimers comprising a sensor domain that detects their cognate stimuli, a catalytic ATP binding domain (CA) that binds ATP and upon signal detection, autophosphorylates a conserved histidine in the dimerization and histidine phosphotransfer domain (DHp). Subsequently, the phosphate is shuttled to a conserved aspartic acid in the response regulator (RR), typically leading to dimerization and high affinity binding to regulatory motifs in the genome(13, 14). The CA and DHp domains are well conserved in HKs within and among different bacteria(14), making them good target sites for the simultaneous inhibition of multiple TCSs and the design of broad-spectrum agents. This polypharmacological approach is expected to limit the emergence of target-based resistance mechanisms(15). Due to the essentiality of some TCSs (e.g. WalKR in *Staphylococcus aureus*(16, 17) and *Bacillus subtilis*(18)) and their role in adaptation, it has been proposed that inhibition of multiple TCSs would compromise growth and/or attenuate survival and virulence in the host(17, 19).

One approach for inhibition of multiple TCSs has been the design of ATP-competitive inhibitors of HKs, with the aim of inhibiting autophosphorylation and TCS signalling(19). The ATP-binding pocket of HKs adopts the Bergerat fold, an α/β sandwich characterized by four conserved regions (the N-box, the G1-box, the G2-box and the G3-box) and the highly variable ATP-lid(20). The Bergerat fold has been observed in the ATP-binding domain of the GHKL protein superfamily, which includes DNA gyrases, the molecular chaperone Hsp90, bacterial HKs and the MutL mismatch repair enzyme(20). Due to this similarity, ATP-competitive Hsp90 inhibitors were proposed as hits for HK inhibition(21). Multiple inhibitors targeting the ATP binding pocket of Hsp90 have been described(22, 23) due to its attractiveness as a target for anti-cancer drugs(24) and some have been shown to weakly bind to the HK PhoQ from *Salmonella*(21). Vo *et al.*(25), showed that among six well-established Hsp90 inhibitors, CCT018159(26), a 3,4-diphenylpyrazole (Figure 1A), was the most potent HK inhibitor of CckA from *Caulobacter crescentus* (IC_50_ = 30 µM) and PhoQ from *Salmonella typhimurium* (IC_50_ = 261 µM). Structure-activity relationship (SAR) investigation around the diphenylpyrazole scaffold led to marginal improvements in HK inhibition, with compound **5b** representing the most potent inhibitor of the series (Figure 1A, IC_50_ CckA = 14 µM; IC_50_ PhoQ = 238 µM). Compound **5b** also showed moderate antibacterial activity against *E. coli* DC2 (a hypersensitive *E. coli* strain), *C. crescentus* and *B. subtilis* (MIC range 12-74 μg/ml)(25).

**Figure 1.**
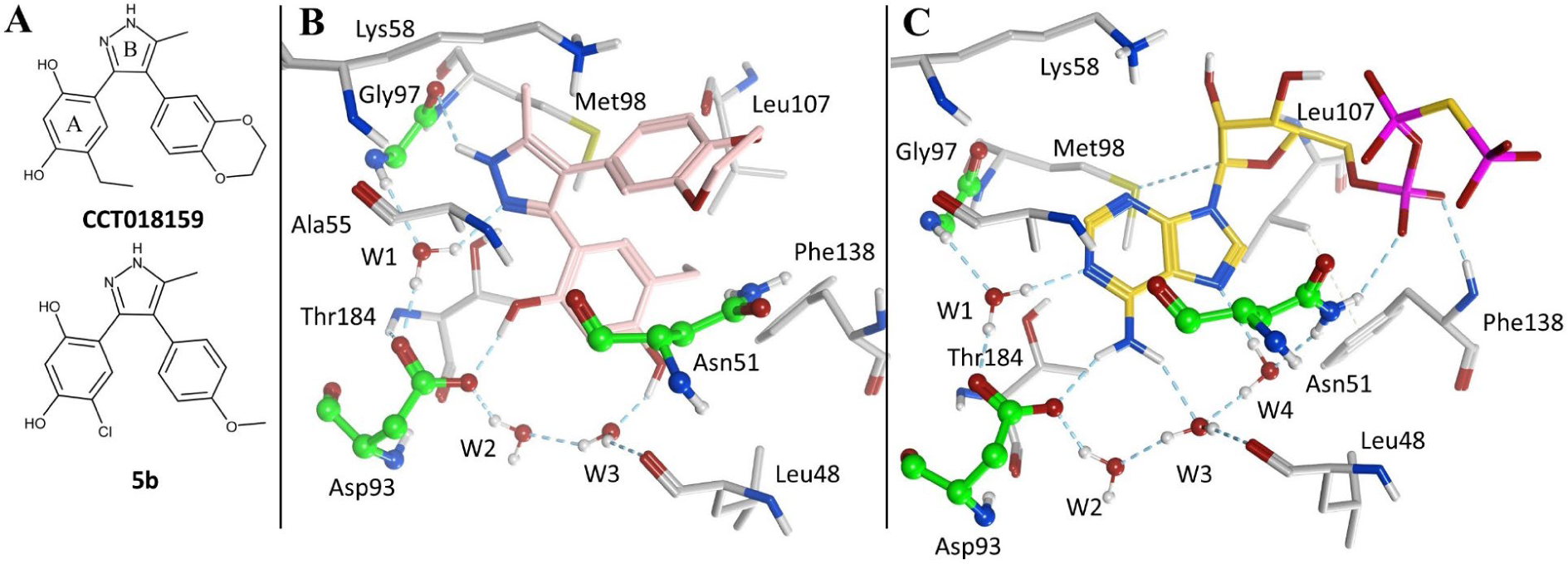
Reported DPP-based HK inhibitors and experimental binding mode to Hsp90α. **A.** 2D chemical structure of representative HK inhibitors with the DPP scaffold that includes the resorcinol (ring A) and pyrazole systems (ring B). **B.** Crystal binding mode of CCT018159 (pink carbon atoms) in complex with Hsp90α (PDB 2BT0(27)). **C.** Crystal binding mode of a non-hydrolyzable ATP analogue (phosphomethylphosphonic acid adenylate ester; yellow carbon atoms) in complex with Hsp90α (PDB 3T10(28)). The carbon atoms of residues highly conserved in HKs are depicted in green while remaining residues in contact with the ligand are coloured in grey. Hydrogen bonds are shown as dashed lines. Image generated in MOE(29).

The 3,4-diphenylpyrazoles are anchored to the ATP-binding site of Hsp90 through a network of hydrogen bonds established by the resorcinol and the pyrazole systems (rings A and B respectively, Figure 1A). As illustrated by the CCT018159-Hsp90α crystal complex in Figure 1B (PDB 2BT0(27)), the 1-hydroxyl from the resorcinol group forms a direct hydrogen bond with the side chain of a buried aspartate residue (Asp93) while the 3-hydroxyl establishes water-mediated interactions. The pyrazole’s N2 forms a water-bridged hydrogen bond with Asp93 and the backbone NH of Gly97. The residues Asn51, Asp93 and Gly97, together with the ordered water molecules (W1, W2 and W3), are highly conserved in HKs as they are involved in binding the adenine system of ATP (Figure 1C). Hence, interactions with these residues are likely to be preserved when repurposing Hsp90 inhibitors as HK inhibitors. The potential for off-target cytotoxic effects by inhibition of mammalian Hsp90 have to be taken into account when using the repurposing approach.

In a previous study, we have reported synthetic derivatives featuring a 3,4-diphenylpyrazole core with antibacterial activity against *S. aureus* (30). Herein, we investigate the antibacterial activity and potential inhibition of HKs for a subset of these derivatives and related compounds from commercial vendors as well as the mechanisms responsible for their antibacterial activity. By studying possible off-target effects and cytotoxicity in mammalian cancer cells, we showed that 3,4-diphenylpyrazoles were inhibiting bacterial growth by interfering with membrane integrity while retaining cytotoxicity for mammalian cells via different mechanisms, membrane damage and Hsp90 inhibition. This highlights the challenges of repurposing Hsp90 inhibitors as antimicrobials targeting TCS HKs but indicates possibilities to exploit 3,4-diphenylpyrazole analogues as agents specifically targeting bacterial membranes. Finally, X-ray crystal structure of resorcinol bound to HK CheA looked attractive for the fragment-based design of novel HK inhibitors.

## Results

### Structure-activity relationship (SAR) of antimicrobial 3,4-diphenylpyrazole (DPP) compounds against *S. aureus*

A collection of 24 DPP analogues was synthesized(30) or purchased to further expand the SAR around the diphenylpyrazole scaffold (Table 1). These analogues were first tested for minimal inhibitory concentration (MIC) against the Gram-positive *S. aureus*.

**Table 1.** Structure, activity against *S. aureus* and toxicity in HEK cells of a series of diphenylpyrazole compounds. Activity against *S. aureus* is expressed as minimal inhibitory concentration in micrograms per milliliter. Toxicity in HEK cells is expressed as LC_50_ (concentration required to kill of 50% of the cell population). Compound substitutions (R1 to R5 and X) from the A, B or C rings of the scaffold as indicated in the diagram are also depicted.

| DPP comp. | X | R<sub>1</sub> | R<sub>2</sub> | R<sub>3</sub> | R<sub>4</sub> | R<sub>5</sub> | MIC *S. aureus*<br>μg/ml (μM) | LC<sub>50</sub> HEK293<br>cells<br>μg/ml (μM) |
| --- | --- | --- | --- | --- | --- | --- | --- | --- |
| 1 | N | -CF<sub>3</sub> | -OH | -H | -H | -H | 6.25 (19.53) | 4.06 (12.69) |
| 2 | N | -CF<sub>3</sub> | -OMe | -H | -H | -H | 12.50 (37.3)† | 7.34 (21.91) |
| 3 | N | -CF<sub>3</sub> | -OEt | -H | -H | -H | 6.25 (17.86) | 25.64 (76.53) |
| 4 | N | -CF<sub>3</sub> | -OBn | -H | -H | -H | 1.56 (3.80)† | 7.26 (17.71) |
| 5 | N | -CF<sub>3</sub> | -OH | -H | -Cl | -H | 3.12 (8.79) | 12.88 (36.28) |
| 6 | N | -CF<sub>3</sub> | -OMe | -H | -Cl | -H | 3.12 (8.43)† | 14.30 (38.65) |
| 7 | N | -CF<sub>3</sub> | -OEt | -H | -Cl | -H | 3.12 (8.21) | 10.82 (28.47) |
| 8 | N | -CF<sub>3</sub> | -OBn | -H | -Cl | -H | 1.56 (3.51)† | 14.25 (32) |
| 9 | N | -Me | -OH | -H | -Cl | -H | 25 (83.33)† | 12.30 (41) |
| 10 | N | -Me | -OMe | -H | -Cl | -H | 25 (79.37) | 23.23 (73.74) |
| 11 | O | -CF<sub>3</sub> | -OH | -H | -Cl | -H | 3.12 (8.76)† | 3.41 (9.61) |
| 12 | N | -CF<sub>3</sub> | -OH | -H | -OMe | -H | 12.50 (35.71) | 31.74 (90.69) |
| 13 | N | -CF<sub>3</sub> | -OMe | -H | -OMe | -H | 12.50 (34.34)† | 29.88 (82.09) |
| 14 | N | -CF<sub>3</sub> | -OEt | -H | -OMe | -H | 6.25 (16.45) | 6.57 (17.29) |
| 15 | N | -CF<sub>3</sub> | -OBn | -H | -OMe | -H | 3.12 (7.09) | 9.64 (21.91) |
| 16 | N | -CF<sub>3</sub> | -OH | -OMe | -H | -H | 25 (71.43) | 16.35 (46.71) |
| 17 | N | -CF<sub>3</sub> | -OMe | -OMe | -H | -H | 25 (68.49) | 12.58 (34.47) |
| 18 | N | -CF<sub>3</sub> | -OH | -H | -OMe | -OMe | 50 (131.58) | 22.81 (60.03) |
| 19 | N | -CF<sub>3</sub> | -OMe | -H | -OMe | -OMe | 25 (63.29) | 13.65 (34.56) |
| 20 | N | -CF<sub>3</sub> | -OEt | -H | -OMe | -OMe | 12.5 (30.49) | 16.53 (40.32) |
| 21* | N | -CF<sub>3</sub> | -H | -H | -Cl | -H | 1.56 (4.8)† | 7.12 (21.91) |
| 22 | N | -CF<sub>3</sub> | -H | -H | -Cl | -H | 1.56 (4.59) | 2.89 (8.5) |
| 23** | N | -CF<sub>3</sub> | -OH | -H | -H | -H | 250 (961.54)† | >250 (>650) |
| 24** | N | -CF<sub>3</sub> | -OMe | -H | -H | -H | 25 (102.04)† | 6.79 (27.71) |

\* 1-OH is absent \*\* C-ring is absent † Data previously published in Solomin *et al*(30).

With the exception of **DPP-23**, all the compounds inhibited *S. aureus* growth with MIC between 1.56 and 50 μg/ml (Table 1). Differently to the parent Hsp90 inhibitor series, several analogues tested are O-alkylated at the R2 positions of ring A, the antibacterial potency of the methoxy and ethoxy analogues is generally comparable to the hydroxyl derivatives, whilst the benzyloxy are among the most potent compounds. Interestingly, removing one (**DPP-22**) or both hydroxyl groups (**DPP-21**) of the resorcinol system results in highly active compounds.

In ring B, the isoxazole acts as a pyrazole bioisostere as it does not alter the MIC (compare **DPP-5** with **DPP-11**). Replacing the R_1_ methyl with a trifluoromethyl group (see pairs **DPP-5**/**DPP-9** and **DPP-6**/**DPP-10**) increases antibacterial potency. Removal of ring C is well tolerated in the R2-methoxy derivative **DPP-24** but significantly decreases the MIC in the resorcinol analogue **DPP-23**. When ring C is present, the R4-chloro derivatives are more active against *S. aureus* than matched compounds lacking the chlorine substituent.

All tested compounds were found to be toxic to the cancer cell-line HEK293 (Table 1). However, only a moderate correlation was found between MIC and toxicity (Spearson correlation r = 0.4289, p-value = 0.0411, Suplementary Figure SF1 A), indicating that mechanisms of antimicrobial activity and toxicity may be different.

### Spectrum of activity

To determine the potential of the DPP compounds as broad-spectrum antibiotics, we tested their antibacterial activity against the Gram-positive organisms *E. faecium* and *E. faecalis*, as well as the Gram-negatives *E. coli, P. aeruginosa* and *P. haemolytica,* which are all important human or animal pathogens (Table 2 and supplementary data, Table ST1). The range of activity of the compounds is similar in all Gram-positive bacteria. The tested compounds show no activity against Gram-negative *E. coli* and *P. aeruginosa* and only moderate activity against *P. haemolytica*.

**Table 2.** MIC data for relevant DPP compounds tested against a panel of Gram-positive and Gram-negative strains, including *E. coli* outer membrane and efflux mutants.

| DPP<br>comp. | MIC (µg/ml) |  |  |  |  |  |  |
| --- | --- | --- | --- | --- | --- | --- | --- |
|  | *E. faecium* | *E. faecalis* | *P.*<br>*haemolytica* | *P.*<br>*aeruginosa* | *E. coli* | *E. coli*<br>JW5503 | *E. coli*<br>D21f2 |
| 2 | 25 | 25 | 50 | >250 | >250 | 6.25 | 50 |
| 5 | 12.5 | 12.5 | 12.5 | >250 | 50 | 3.12 | 50 |
| 6 | 6.25 | 6.25 | 12.5 | >250 | >250 | 3.12 | 25 |
| 7 | 3.12 | 3.12 | 25 | >250 | >250 | 3.12 | 25 |
| 12 | 50 | 50 | 50 | >250 | >250 | 12.5 | >250 |
| 13 | 25 | 25 | >250 | >250 | >250 | 12.5 | >250 |

We also tested the inhibitory activity of compounds in two different *E. coli* mutants: i) *E. coli* JW5503 that lacks TolC efflux pumps and ii) *E. coli* D21f2 that has a defective LPS inner core increasing permeability of the outer membrane. *E. coli* D21f2 has a slightly higher susceptibility to DPP compounds than *E. coli* ATCC25922. MIC data for *E. coli* JW5503 is similar to that obtained for Gram-positive bacteria, indicating that TolC-dependent efflux plays a major role in the susceptibility of *E. coli* to the DPP inhibitors. Table 2 shows MIC data of the most relevant compounds in the panel of strains tested.

### Docking and inhibition studies

Docking studies with the HK PhoQ from *E. coli* suggest that only a subset of the DPP compounds can adopt a binding mode similar to the CCT018159-Hsp90α complex, exemplified by the putative binding pose of **DPP-5** (Figure 2 A and B). In detail, the 5-membered ring of **DPP-5** forms water-mediated contacts with Asp415 and Gly419, whilst Ile420 and Tyr393 flank the two faces of the pyrazole. Asp415 and Gly419 are part of the G1-box and are highly conserved among members of the GHKL family (Figure 2C). On the contrary, aromatic residues at the position corresponding to Tyr393 position are HK-specific (this residue corresponds to Ala55 in Hsp90α, Figure 2C)(20). Ring C and the trifluoromethyl group are projected towards the opening of the ATP-binding pocket. The 1-hydroxyl of the resorcinol scaffold engages the conserved aspartate (Asp415), whilst the 3-hydroxyl binds the carbonyl backbone of Val386. However, by analogy with the CCT018159 binding mode in Hsp90, the 3-hydroxyl could also participate in the hydrogen bond network with a conserved water molecule system (W2-W3). The docking settings did not include the water molecules W2 and W3 to allow for their potential displacement when the 3-hydroxy is alkylated. Due to the limited size of the subpocket housing these water molecules (shown as a surface in Figure 2), derivatives with bulky substituents in position 3, such as the benzyloxy group of **DPP-8**, are unlikely to retain this binding mode unless there are major protein conformational changes. The *in silico* studies predict a flipped binding mode for **DPP-8**, in which the pyrazole nitrogen atoms are hydrogen bonded to Asp415 and W1, and the trifluoromethyl group displaces the water network. The ring A is flanked by Gly419, Ile420 and Tyr393, with the benzyloxy group partially solvent exposed, establishing contacts with Pro418 and Pro421. Ring C faces the hydrophobic floor of the pocket formed by the side chains of Ile428, Leu446 and Met472. An analogous binding pose is also predicted for the derivative lacking both hydroxyl groups (**DPP-22**).

**Figure 2.**
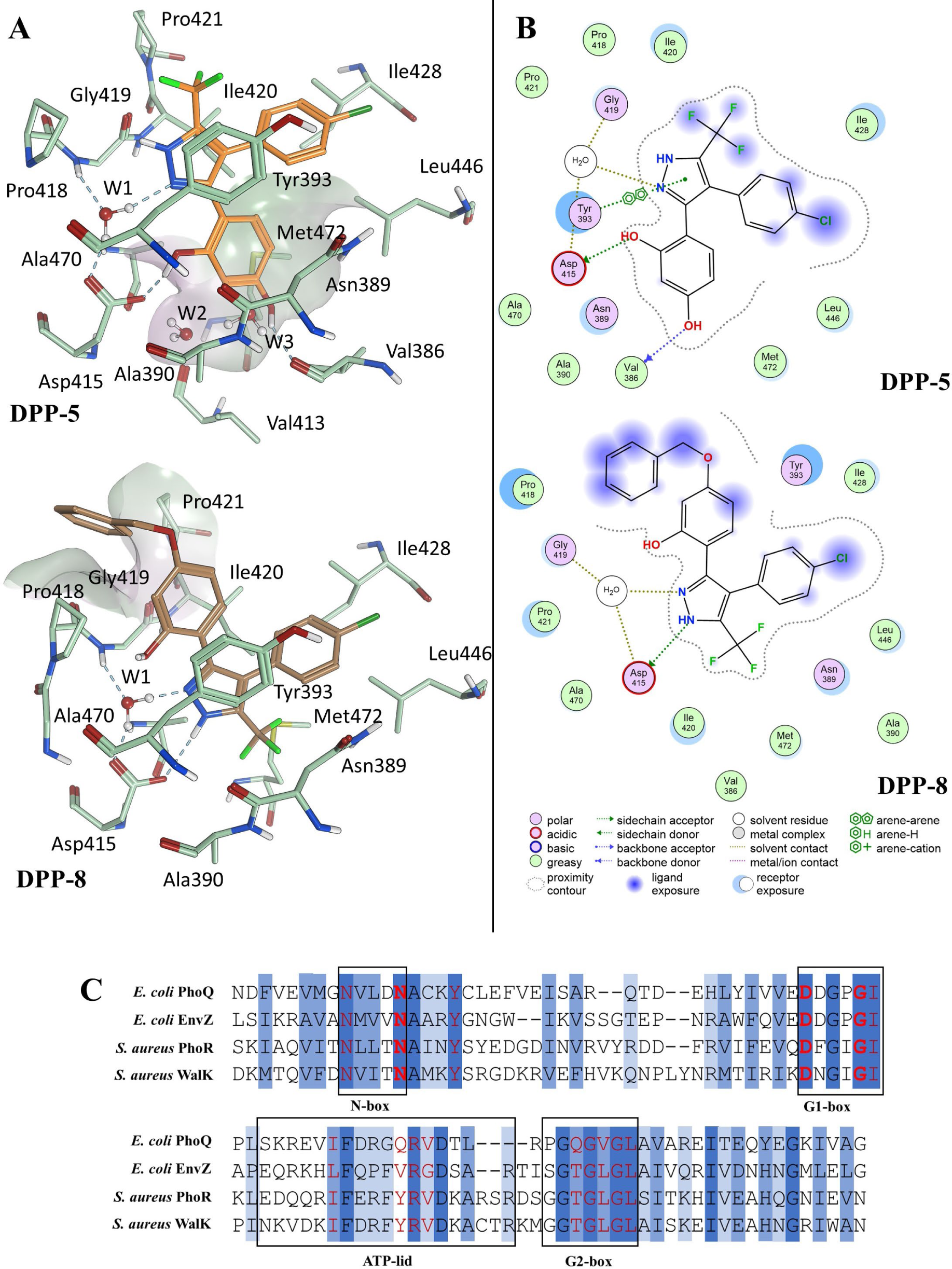
*In silico* binding modes of representative DPP compounds and HK sequence alignment. **A)** 3D models of the top ranked GOLD docking pose of **DPP-5** and **DPP-8** in complex with the HK PhoQ from *E. coli* (PDB 1ID0). A surface delineates: the subpocket housing W2-W3 in the **DPP-5** pose; the putative interaction site of the benzyloxy of **DPP-8** (Surface colour code: purple = hydrophobic; white = Neutral; Green = Lipophilic). Image prepared using MOE. **B)** MOE ligand interaction diagrams of the top ranked GOLD docking pose of **DPP-5** and **DPP-8** in complex with the HK PhoQ from *E. coli* (PDB 1ID0). **C)** Protein sequence alignment of *E. coli* PhoQ, *E. coli* EnvZ, *S. aureus* PhoR and *S. aureus* WalK. Amino-acids in red are predicted to interact with ATP. In red and bold the conserved triad (N389, D415 and G419 in *E. coli* PhoQ), Y393 is conserved in HKs but is absent in Hsp90 (dark blue – 100% amino-acid conservation, blue – amino acid physicochemical property conserved, light blue – amino acid conserved in at least 80% of the HKs).

The compounds were tested for inhibition of HK autophosphorylation *in vitro.* Of the 24 derivatives, 6 inhibited *in vitro* autophosphorylation of *E. coli* EnvZ at a concentration of 2 mM. For 4 compounds that showed total inhibition in the initial screening, the IC_50_ (amount of compound required to reduce HK autophosphorylation by 50%) was measured and was in the high micromolar range for *S. aureus* PhoR and *E. coli* EnvZ (Table 3 and supplementary data, Figure SF2). Each compound has similar IC_50_ values for PhoR and EnvZ, indicating good potential for polypharmacology. We discounted the notion that DPP-induced aggregation of HKs was the reason for inhibiting autophosphorylation using native gel protein electrophoresis of HK proteins with and without DPPs (data not shown). We also investigated if the four DPP inhibiting HK autophosphorylation would also bind to Hsp90α in a fluorescence polarization assay. All the compounds tested displaced FITC-labelled geldanamycin with IC_50_s in the nM range (Table 3).

**Table 3.** *In vitro* inhibition of HK autophosphorylation by DPP compounds and Hsp90 fluorescence polarization assay results. Six DPPs showed inhibition of EnvZ autophosphorylation *in vitro.* The IC_50_ ± SD (concentration of inhibitor that reduces HK autophosphorylation by 50% ± standard deviation) was calculated for 5 of these DPPs using *S. aureus* PhoR and *E. coli* EnvZ (n = 2). The results of the Hsp90 fluorescence polarization assay are reported as IC_50_s (compound concentration that decreases geldanmycin binding by 50%)

| Compound | $IC_{50}$ PhoR<br>*S. aureus* ($\mu M$) | $IC_{50}$ EnvZ<br>*E. coli* ($\mu M$) | $IC_{50}$ Hsp90<br>($\mu M$) |
| --- | --- | --- | --- |
| DPP-2 | n.t. | <2 mM | n.t. |
| DPP-5 | $95 \pm 5.6$ | $117 \pm 6.3$ | 0.496 |
| DPP-6 | $55 \pm 3.8$ | $98 \pm 11.84$ | 0.798 |
| DPP-7 | n.t. | <2 mM | n.t. |
| DPP-12 | $328 \pm 4.1$ | $427 \pm 9.1$ | 0.574 |
| DPP-13 | $89 \pm 6.9$ | $175 \pm 10.1$ | 0.838 |

We also tested whether the DPP series would inhibit the bacterial gyrase, which also possesses the Bergerat fold in their ATP-binding domain(20), as this would lead to antimicrobial activity. None of the DPP compounds inhibited gyrase activity, indicating this mechanism of action is not responsible for their antimicrobial activity (supplementary data, Figure SF3).

### Resorcinol fragment binds to the ATP-binding site of the histidine kinase CheA

Our attempts to co-crystallize the DPP derivative with the CA domain of CheA from *T. maritima*, a class II HK for which robust crystallization protocols have been established, failed. However, we successfully obtained the X-ray crystal structure of the resorcinol fragment bound to CheA-CA with a resolution of 2.1 Å. The binding mode of the resorcinol resembles the predicted binding mode of **DPP-5** (Figure 2) and is nearly identical to the interaction formed by CCT018159 with Hsp90α (Figure 1). The overall structure showed a solvent-accessible binding pocket where the resorcinol was observed making contact with the conserved aspartate (Asp449) (Figure 3 A and B): the one hydroxyl from the resorcinol is in the proximity of the conserved aspartate (Asp449) and the ordered water molecule W1, while the other hydroxyl of the resorcinol forms water-bridged contacts with the backbone carbonyl a leucine residue (Leu406; Val386 in PhoQ *E. coli*). The crystal binding mode of the resorcinol fragment supports the hypothesis that resorcinol-containing DPP compounds are likely to retain the binding mode observed in Hsp90.

**Figure 3.**
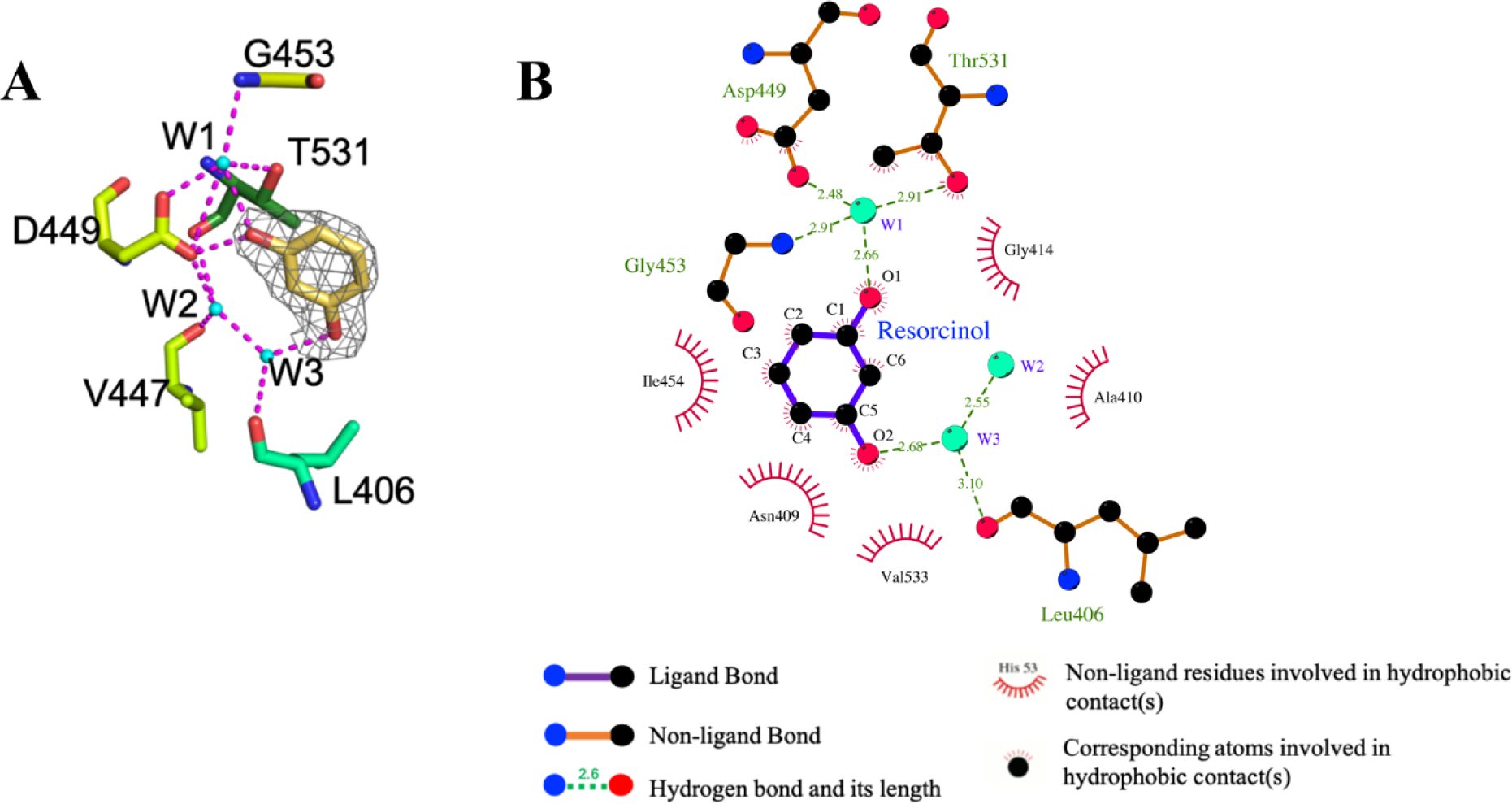
Resorcinol binding mode to the binding pocket of CheA-CA domain. A. Fo-Fc map shows the electron density around the resorcinol (resolution 2.1 Å). Image prepared using PyMol B. Binding mode prediction of resorcinol to CheA-CA domain, created in LigPlot^+^(31). Resorcinol makes direct and indirect contacts through water molecules (W1 and W2) with the conserved Aspartate (D449) residue of CheA.

### DPPs increase membrane permeability in *S. aureus*

To study the effect of DPP compounds on the cytoplasmic membrane of *S. aureus* we used the nucleic acid stain SYTOX Green. DPPs were tested for loss of membrane integrity after 5 minutes exposure to 0.5x, 1x and 2xMIC. The tested DPP compounds significantly increased membrane permeability, similar to nisin which was used as positive control (Figure 4, supplementary Table ST2). In contrast, no effect on membrane integrity was observed with the gyrase inhibitor novobiocin at 2xMIC. **DPP-4**, **DPP-5** and **DPP-15** were the only compounds that did not cause significant loss of membrane integrity at 1xMIC.

**Figure 4.**
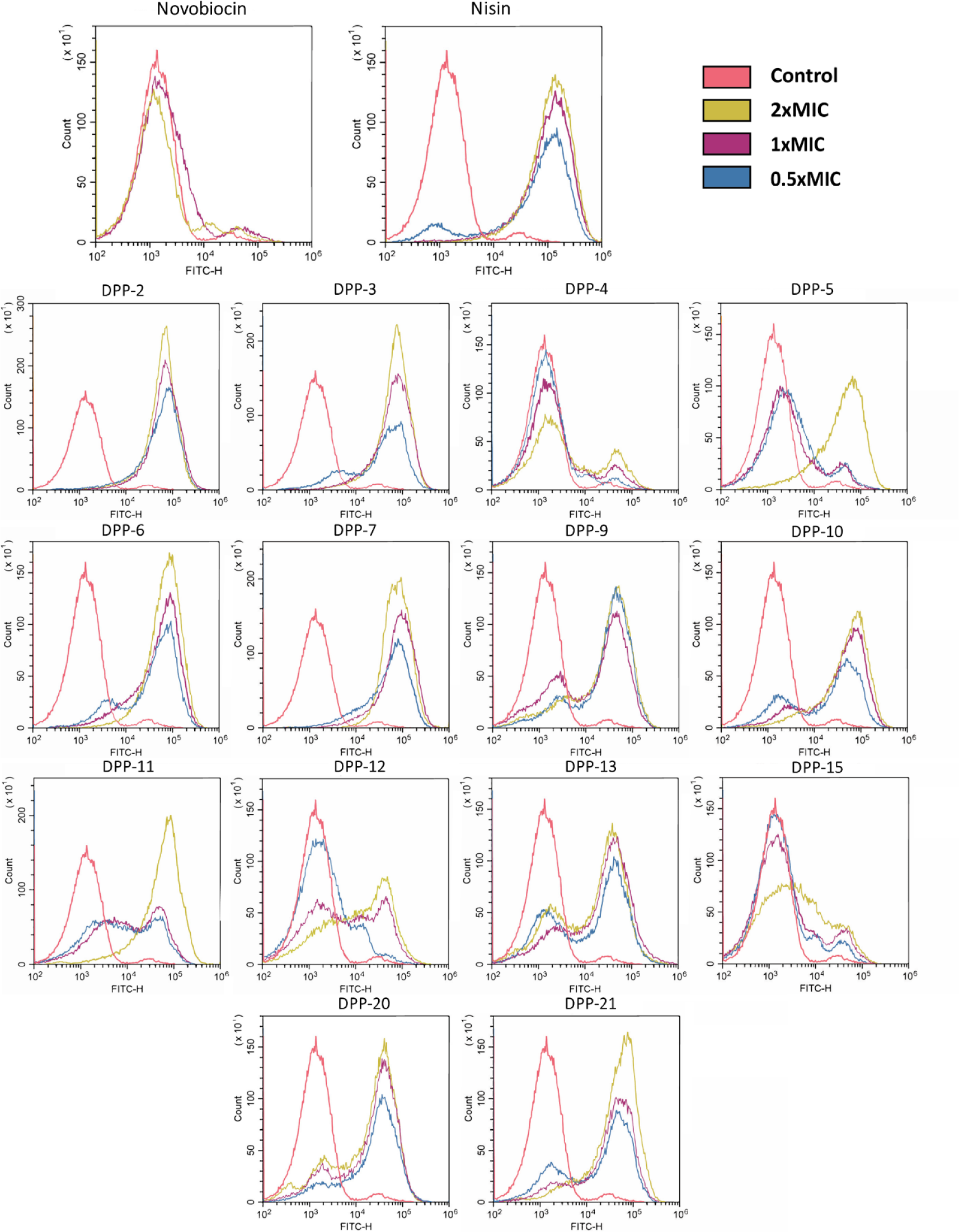
Permeabilization of *S. aureus* membrane by DPP compounds. *S. aureus* membrane permeabilization was measured using SYTOX-Green fluorescent stain, which can bind to nucleic acid if it penetrates cytoplasmic membranes. Permeabilization is measured as the shift in FITC-H fluorescence compared to the non-treated control (red). DPPs were tested at 0.5 (blue), 1 (purple) and 2xMIC (yellow) with an incubation time of 5 minutes. Nisin was used as positive control for permeabilization and novobiocin as a negative control.

To see if the membrane destabilizing activity of DPP compounds was specific for bacterial membranes, we tested their haemolytic activity with sheep red blood cells (RBCs) (Figure 5). There was no correlation between the percentage of RBC haemolysis and the membrane damage caused in *S. aureus* (Spearson correlation r = 0.4402, p-value = 0.1152, Suplementary Figure SF1 B).

**Figure 5.**
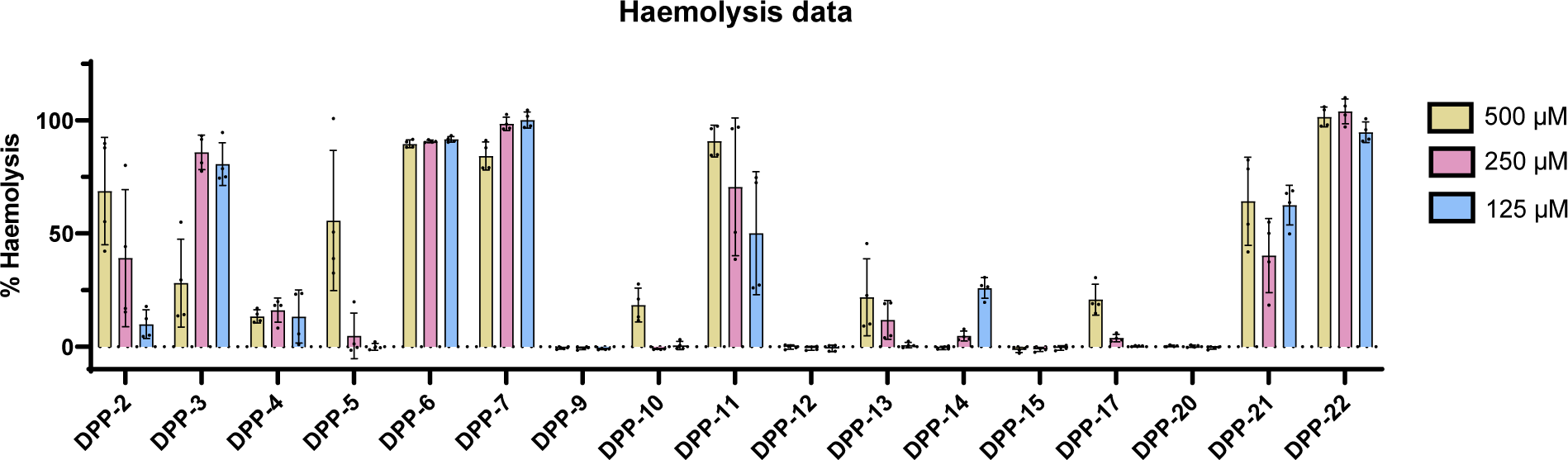
Haemolysis of sheep red blood cells by DPPs. Percentage of RBC haemolysis compared to Triton-X control is shown after 30 min exposure to 500, 250 and 125 μM of each compound; Error bars represent Standard Deviation (n = 4).

### Cytotoxicity of DPPs

The cytotoxicity of DPP compounds was quantified using the Alamar Blue assay and the human embryonic kidney cell line HEK293. All compounds with MIC ≤ 50 μg/ml against *S. aureus* were cytotoxic, with LC_50_ values ranging from 2.89 to 31.7 μg/ml (Table 1). Eight out of 20 DPPs tested caused significant haemolysis of sheep RBCs at 250 μM after 30 min incubation (Figure 5). We did not see a significant correlation between haemolysis and cytotoxicity LC_50_s (Spearson correlation r = -0.351, p-value = 0.1671, Suplementary Figure SF1 C). Since we confirmed that the DPP compounds bind to Hsp90, we hypothesized that there may be two different mechanisms of cytotoxicity, with some DPP analogues inducing membrane-associated damage to mammalian cells (such as **DPP-6**), and others through Hsp90 inhibition.

To test this hypothesis, we performed histology on HEK293 cells treated with three cytotoxic DPP compounds: **DPP-6** (that causes 100% RBC haemolysis), **DPP-14** and **DPP-20** (not haemolytic). We did not observe membrane blebbing or other morphologies associated with membrane damage(32) in cells treated with **DPP-6** or the DMSO control (Figure 6B and C). When cells were treated with **DPP-14** (data not shown) and **DPP-20** (Figure 6D), we observed a higher number of cells in G2/M phase of cell cycle (12.24±1.23% and 26.25±4.68% of cells in G2/M phase in **DPP-14** and **DPP-20** respectively compared to the 2.68±1.31% in DMSO control and 1.63±1.10% in media control) which has been reported for Hsp90 inhibitors(33, 34).

**Figure 6.**
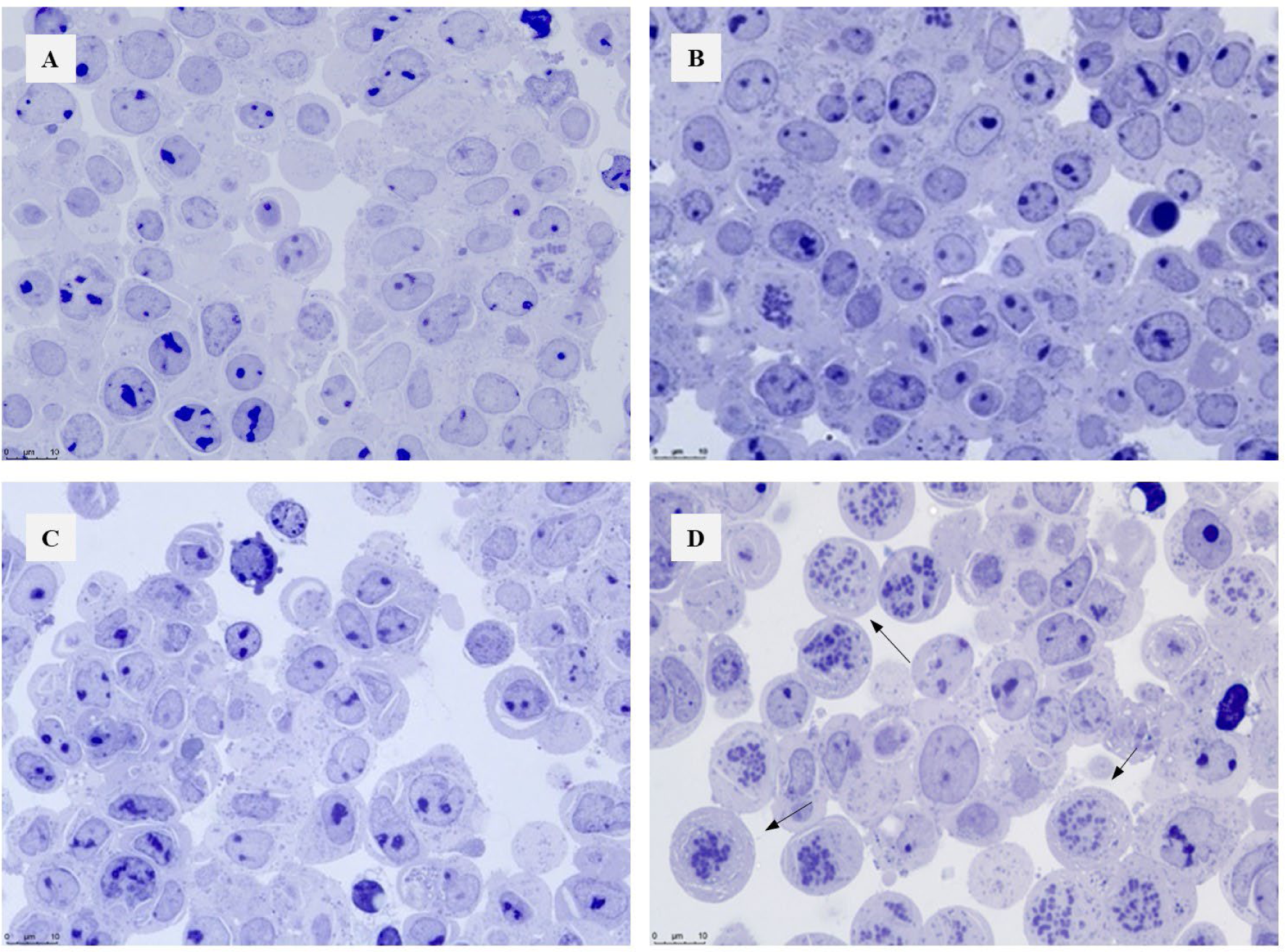
Histological images of HEK293 cells after 24 hours exposure to DPP compounds. A. HEK293 cells grown in exposure media or B. 1% DMSO vehicle control C. HEK293 cells grown in the presence of LC_20_ concentration (2.5 μg/ml) **DPP-6**. D. HEK293 cells grown in the presence of LC_20_ concentration (3 μg/ml) **DPP-20**. Arrows show high proportion of cells in G2/M phase.

## Discussion

Researchers have been trying for over two decades to find inhibitors of bacterial TCSs(35), but no inhibitor has reached clinical evaluation. Here, we build on previous efforts to repurpose mammalian Hsp90 inhibitors targeting the ATP-binding pocket, which has a similar binding fold to the one found in HKs(25). We further explored the chemical space of the 3,4-diphenylpyrazols, as they have been reported to inhibit HKs. Our collection of DPP analogues were shown to inhibit the growth of Gram-positive bacteria but not Gram-negative, likely due to their capacity to efflux small molecules and affect accumulation. This was supported by our finding that the MIC of an *E. coli* Δ*tolC* mutant was similar to that of *S. aureus.* Six DPPs inhibited HK autophosphorylation *in vitro,* but compounds with good antibacterial activity were inactive in the inhibition assay, suggesting that many of these DPPs inhibit bacterial growth through a mechanism other than HK inhibition.

To explore the causality between HK inhibition and antimicrobial activity we first tested for common off-target effects. Hilliard *et al.* (36) demonstrated that the TCS inhibitors described in the literature at that time interfered with membrane integrity in *S. aureus*, and/or cause haemolysis in mammalian erythrocytes. Most of the tested DPP compounds caused an increase in membrane permeability similar to the nisin control, while 8 out of the 17 compounds tested presented haemolysis in sheep RBC. There was no correlation between haemolysis or mammalian cell death and membrane permeabilizing activity in Gram-positive bacteria, suggesting different mechanisms of membrane disruption in bacteria and mammalian cells. This opens up new possibilities for optimizing these compounds as specific bacterial membrane disruptors rather than TCS inhibitors.

Docking studies indicate that the substitution pattern on ring A influences the binding mode, with the resorcinol-based derivatives likely to adopt a binding mode analogous to the crystal-bound complex of CCT018159-Hsp90α; an observation also supported by the X-ray crystal structure of resorcinol in complex with the HK CheA. Compounds with bulky substitutions such as **DPP-8** or that do not contain the resorcinol ring (**DPP-21**) were predicted to have a flipped pose. None of the compounds predicted to bind in the flipped orientation were found to inhibit HK autophosphorylation *in vitro,* possibly indicating lower affinity for the ATP pocket. Some compounds that were likelty to share a similar binding mode **DPP-5** did not inhibit autophosphorylation. This is possibly due to precipitation of the compounds affecting the inhibition assays.

One of the main drawbacks of repurposing Hsp90 inhibitors is the potential of retaining toxicity to mammalian cells due to their inhibitory effect on Hsp90(37). Even though Hsp90 has been identified as an anti-cancer target due to its greater importance in cancer cell protein homeostasis than in normal cells(24), it still plays an important role in normal metabolism(38). Therefore, when repurposing Hsp90 inhibitors it is important to find chemical spaces that favour binding to HKs while diminishing binding to Hsp90. We saw that DPP compounds still bound with high affinity to Hsp90, thus, more efforts are needed to dissociate Hsp90 activity from HK inhibition.

Finally, we also showed that Hsp90 inhibitors can cause toxicity *in vitro* in HEK293 cells. As some DPPs caused haemolysis, we performed histological analysis to see if lethality in cell lines was produced by plasma membrane disruption or Hsp90 inhibition. We could not see significant differences in the phenotype of cells treated with LC_20_ concentration of the highly haemolytic **DPP-6** compared to the control. However, this could be due to lack of effect at this low concentration. **DPP-20**, a non-haemolytic analogue, caused cell-cycle arrest in G2/M phase as shown for some inhibitors of Hsp90(33, 34). Based on these findings we suggest that cytotoxicity by non-haemolytic DPPs is due to Hsp90 inhibition.

In summary, we further explored the repurposing of the DPP scaffold from inhibition of Hsp90 to inhibition of HKs. Structural analogues were synthesized to explore the chemical space for inhibition of HK autophosphorylation and binding to HKs. Even though in this study we highlight some of the difficulties in using DPP analogues for antimicrobial discovery, we were able to demonstrate different mechanisms of action for inhibition of bacterial growth and cytotoxicity. Optimization of these compounds as potential bacterial membrane disruptors could, together with reducing affinity to Hsp90 lead to good antimicrobials with low toxicity. Finally, the high-resolution crystal structure of the resorcinol fragment can be used as a starting point for the structure-based design of selective HK inhibitors, with compound properties carefully optimized to avoid off-target effects and toxicity in mammalian cells.

## Material and Methods

### Chemical compounds

The chemical compounds **DPP-2**, **4**, **6**, **8**, **9**, **11**, **13**, **21**, **22**, **23** and **24** were synthesized by the Latvian Institute of Organic Synthesis. Details of the synthesis of the compounds can be found in Solomin *et al*.(30); the schematic synthesis of **DPP-22** is shown in Supplementary Figure 4 (Figure SF4). Chemical compounds **DPP-1**, **3**, **5**, **7**, **10**, **12**, **14**, **15**, **16**, **27**, **28**, **19** and **20** were purchased from MolPort (Riga, Latvia). Working stocks of compounds were prepared in dimethyl-sulfoxide (DMSO) at a concentration of 50 mg/ml.

### Bacterial strains and growth conditions

*Staphylococcus aureus* str. Newman, *Enterococcus faecalis* vanA (strain E0155), *Enterococcus faecium* vanA (strain E1654), *Escherichia coli* ATCC 25922, *Escherichia coli* D21f2 with a truncated LPS barrier, *Escherichia coli* JW5503 (Δ*tolC*)(39), *Pseudomonas aeruginosa* ATCC 27853 and *Pasteurella haemolytica* ATCC 29701 were grown in Mueller-Hinton Broth (MHB) (Oxoid, Basingstoke, UK) and incubated at 37°C.

### Minimal inhibitory concentration (MIC)

The minimal inhibitory concentration (MIC) was determined using the microdilution method following guidelines of the European Committee on Antimicrobial Susceptibility Testing (EUCAST)(40). Briefly, a series of two-fold dilutions in MHB of each compound were made in a 96-well plate, with final concentrations range from 50 to 0.39 μg/ml in 100 μl final volume per well. A hundred microliters of 1:100 dilution of an overnight culture of the correspondent bacteria in MHB was added to each well. Plates were incubated for 18 h at 37 °C. MIC was recorded as the lowest concentration where no growth was detected as measured by optical density at 600 nm (OD_600_) using Spectramax M5 (Molecular Devices LLC, San Jose, CA, USA). Compounds that did not inhibit growth were re-tested using a higher concentration range (250-1.95 μg/ml). Wells containing bacteria with or without 1% DMSO and medium alone were included as controls in every plate.

### Cell culture and cytotoxicity using Alamar blue assay

Human embryonic kidney cells (HEK293) cells and the human hepatocellular carcinoma cells (HepG2) were routinely cultured in 75 cm^2^ culture flasks (Corning Incorporated) on high-glucose Dulbecco’s Modified Eagle Medium (DMEM) containing glutaMax and phenol red (Gibco) supplemented with 1% penicillin/streptomycin (Gibco) and 10% fetal bovine serum (FBS) (Gibco). Cells were maintained at 37 °C and 5% CO_2_.

Cytotoxicity assays with HEK293 and HepG2 cells were performed in 96-well plates seeded with 5 x 10^4^ cells/well and incubated for 24 h to reach 80-90% confluency. Exterior wells were filled with only medium to prevent evaporation. Culture medium was removed from the cells and 100 μl of exposure medium (high-glucose DMEM without phenol red (Gibco)) was added to avoid potential interaction between FBS components or antibiotics and tested compounds. A two-fold dilution series of compounds in exposure medium (range 100-3.2 μg/ml) was made in a separate 96-well plate and 100 μl of each of these dilutions was added to the assay plates for 24 hours at 37 °C in presence of 5% CO_2_. Control wells contained cells with DMEM, DMEM+1% DMSO (vehicle control) or DMEM+20% DMSO. Wells without cells were also included as negative control. After exposure, the medium was replaced with 100 μl 10% Alamar Blue (Invitrogen) in exposure medium. After 45 minutes incubation at 37°C and 5% CO_2_, fluorescence was measured on a Spectramax M5 plate reader at λex = 541 nm, λem = 590 nm. Cell viability compared to the vehicle control was calculated and inhibition curves for each compound were fitted (non-linear curve fit, four variables, bottom constrained = 0) in Prism 9 (Graphpad Software, San Diego, USA). The LC_50_ value for 50% cell viability was calculated based on the fitted curves.

### Computational studies

The compounds were prepared for docking using the MOE(29) database wash application to add hydrogens, to generate protonation states and tautomers. The MOE energy minimization application was employed to generate low-energy conformations using the MMFF94x forcefield. The protein with PDB code 1ID0(41) was imported from the RSCB PDB(42). As implemented in MOE, the protein preparation application was used to add hydrogens, assign bond orders, build missing side chains, and assign protonation states. Only chain A was retained for the docking. Water molecule HOH16 (hereafter defined as W1) was retained, while the remaining water molecules and metals were removed. The prepared compounds were docked using the GOLD-5.2(43) molecular docking tool. The binding site was defined by the protein atoms within 9 Å from the crystal-bound ligand. W1 was set as fixed (toggle state: on; spin state: fix). ChemPLP was used as the scoring function. The search efficiency was set to very flexible. A protein HBond constraint to the conserved aspartic acid residues was added to penalize poses not forming such an interaction (constraint weight: 10; minimum H-bond geometry weight: 0.005).

### Protein production and Purification

Cytoplasmic domains from *S. aureus* PhoR and *E. coli* EnvZ as well the CA domain of *Thermotoga maritima* CheA were cloned in pNIC28-Bsa4(44) plasmid containing His_6_-tag were expressed in *E. coli* RIL and purified as previously described(45–47) using His-affinity and size exclusion chromatography. Briefly, *E. coli* RIL strains carrying the appropriate plasmid were grown in 1 l Luria-broth (LB) (Merck Millipore) supplemented with kanamycin (100 μg/ml) with shaking (200 r.p.m.) at 37 °C . When OD_600_ reached 0.5, 1 mM IPTG was added to induce protein expression and cells were grown for 3 more hours. Cells were harvested by centrifugation (4.000 g, 4 °C), resuspended in 40 ml buffer A (50 mM Tris-HCl pH 8.0, 0.5 M NaCl, 10% glycerol and 1 mM phenylmethanesulfonyl fluoride), sonicated (4 °C, 5 min with pulses of 15 sec at intervals of 1 minute) and centrifuged (11.000 g, 4 °C, 60 min). Supernatant was passed through a 5 ml HisTrap HP column (GE Healthcare) equilibrated in buffer A using AKTA system (GE Healthcare). His-column was washed with buffer A (80 ml), and a linear gradient of imidazole (range 0 to 0.25 M) in buffer A was applied. The purest fractions evaluated by SDS-PAGE were collected, concentrated by ultrafiltration in a Amicon Ultra-15 (10 or 30kDa exclusion) (Merck Millipore) and further purified by gel filtration using Superdex 200 column (GE Healthcare), equilibrated in buffer B (50 mM TrisHCl, pH 8.0, 0.2 M NaCl and 5% glycerol). Purified proteins were then concentrated using ultrafiltration and stored at -80 °C.

Hsp90α C-terminal domain was purified as described by Goode *et al*. (48) *E. coli* BL21(DE3) expression strains containing GST-Hsp90 N(9–236) plasmid (Addgene: 22481) were grown in 1 l LB media supplemented with 100 μg/ml ampicillin at 25 °C and shaking (200 r.p.m.) to OD_600_ 0.5 and protein expression was induced by the addition of 1 mM IPTG and the culture was grown at 25 °C and shaking (200 r.p.m.) overnight. Cells were then pelleted (4.000 g, 4 °C), resuspended in 40 ml buffer A2 (50 mM Tris-HCl pH 8.0, 0.5 M NaCl and 1 mM phenylmethanesulfonyl fluoride), sonicated (4 °C, 5 min with pulses of 15 sec at intervals of 1 minute) and centrifuged (11.000 g, 4 °C, 60 min). Supernatant was passed through a 5 ml Glutathione Sepharose 4B (GE Healthcare) equilibrated with buffer A2 using an AKTA system (GE Healthcare). Column was washed using buffer A2 and a linear gradient of glutathione (0-0.25M) in buffer A2 was applied. Fractions containing Hsp90α were collected, mixed and concentrated by ultrafiltration.

### HK autophosphorylation

Autophosphorylation inhibition assays were performed in kinase assay buffer (100 mM Tris-HCl (pH 8.0), 5 mM MgCl_2_, 10 mM DTT), containing 5 μg of purified HK and 10 mM ATP for 30 min at 25 °C. The reaction was stopped by adding gel loading buffer containing 16% SDS before loading the reaction on a 12% SDS-PAGE.. Non-hydrolyzable AMP-PNP (10 mM) was used as control for inhibition of autophosphorylation. In the first screening inhibitors were added at a concentration 2 mM and compounds giving complete inhibition of autophosphorylation were selected for screening at a range of concentrations (0-1000 µM). To detect autophosphorylation, the gel was transferered to a nitrocellulose membrane (GE Healthcare) at 100 V in the transfer buffer(25 mM Tris, 192 mM Glycine, 10% Methanol). The membrane was then blocked using Tris-buffered saline solution with 0.1% v/v Tween 20 (TBS-T) supplied with 5% Bovine serum albumin. The phosphorylated histidine was detected by Western blotting by then incubating the membrane for 60 min with 1:1.000 anti-N1-phosphohistidine antibody (Merck Millipore) in TBS-T followed by a washing step for 30 min with TBS-T and a final 60 min incubation with with 1:10.000 horseradish peroxide-conjugated anti-rabbit IgG (Promega) in TBS-T. Chemiluminescent reaction was measured using Pierce ECL immunoblotting substrate (ThermoScientific) in a LAS 3000 image analyzer (Fujifilm Life Science, Tokyo, Japan) followed by densitometric analysis using Multi Gauge image-analysis software (Fujifilm Life Science, Tokyo, Japan). The inhibitor concentrations required to halve the chemiluminescence intensity (IC_50_) were determined using Prism 4.1 (GraphPad Software, San Diego, CA, USA).

### Hsp90 inhibition using fluorescence polarization

Competitive binding of inhibitors to Hsp90 was measured by fluorescence polarization assay as previously reported(49). Inhibitors were assayed in different concentrations (range 0-1000 µM) in a Nunc black, low binding 384-well plate (ThermoScientific). Each reaction was prepared in reaction mix (100 mM Tris-Cl, pH 7.4, 20 mM KCl, 6 mM MgCl_2_, 2 mM DTT and 0.1 mg/ml bovine serum albumin) in a final volume of 50 μl containing 350 nM Hsp90 and 100 nM geldanamycin FITC-labelled (BPS Bioscience, San Diego, CA, USA), and the inhibitor in the correspondent concentration. Blank control containing no enzyme and no geldanamycin, enzyme positive control containing no inhibitor, and enzyme negative control containing no Hsp90 were included. Fluorescence polarization was then measured using Tecan SPARK (Tecan, Grödig, Austria) at wavelengths λex = 485 nm and λem = 530 nm. Binding of FITC-labelled geldanamycin to Hsp90 results in low fluorescence polarization, competitive binding of the inhibitors resulted in an increase of fluorescence polarization from the free FITC-labelled geldanamycin. IC_50_ values that decrease geldanamycin binding in 50% were calculated using Prism 4.1 (GraphPad Software, San Diego, CA, USA).

### Crystallization and structure determination of the CheA-Resorcinol complex

The CheA-CA domain was crystallized by the sitting-drop vapor-diffusion method at 21°C using 0.4 µL of 25 mg/mL CheA-CA protein mixed with 0.4 µL of reservoir solution containing 30% (w/v) PEG 8000, 0.6 M ammonium acetate, 0.065 M sodium acetate pH 4.5. Monoclinic crystals appeared within 24 hrs and 0.2 µL of 20 mM Resorcinol solution (Enamine, Frankfurt, Germany) was added to the crystallization drops. After 1 hour the crystals were harvested, soaked in 40 % (w/v) PEG 8000 as cryoprotectant and flash frozen in liquid nitrogen. The crystals were diffracted at XALOC beamline ALBA synchrotron, Barcelona at 100 K. Programmes within CCP4 suit were used for data processing(50). The structure was solved by molecular replacement using the published structure (PDB 1I58(51)) as template using PHASER (52) followed by refinement using REFMAC5(53). Automated model building was performed using BUCCANEER (54) followed by manual model building in COOT. The coordinates and restraints for resorcinol were generated using AceDRG(55). COOT was used again for ligand fitting and chain tracing followed by refinement using REFMAC5. The part of the lid loop corresponding to the residues 492-505 could not be built owing to the absence of the corresponding electron density in the area. The data collection and refinement statistics can be found in supplementary data (Table ST3). The coordinates for the solved structure are deposited in the Protein Data Bank with accession code 8PF2.

### Membrane permeability using SYTOX-Green

*S. aureus* str. Newman was grown to exponential phase (OD_600_ = 0.4) in MHB media. Five hundred microliters of bacteria were transferred to 1.5 ml tubes (Eppendorf) and incubated for 5 min with different compounds at concentrations of 0.5xMIC, 1xMIC and 2xMIC. Nisin (Sigma-Aldrich) and novobiocin (Sigma-Aldrich) were used as positive and negative controls for membrane permeabilization, and stained and unstained untreated controls were also included in each assay. After exposure, samples were centrifuged 4.000 r.p.m. for 5 min and resuspended in PBS. Bacteria were dyed by adding the fluorescent nuclear stain SYTOX® Green (Molecular Probes, Invitrogen) to a final concentration of 0.5 μM and incubating for 5 minutes in the dark. Samples were centrifuged again and resuspended with PBS to remove unbound stain. Three hundred microliters of each sample were transferred to a clean well in a 96-well clear-bottom plate (Corning Incorporated). Fluorescence was measured in CytoFLEX® flow cytometer (Beckman Coulter, Brea, CA, USA) using the green (FITC) channel with the excitation wavelength at 488 nm. Overlay histograms were created with CytExpert Software (Beckman Coulter, Brea, CA, USA).

### Haemolysis assays

The haemolysis assay was adapted from Evans *et al* (56). Briefly, sheep red blood cells (RBC) 10% washed pooled cells in PBS (Rockland Immunochemicals, Limerick, PA, USA) were diluted 1:5 to give a final concentration of 2% RBCs in PBS. Two hundred microlitres of the RBC suspension was transferred to 1.5 ml Eppendorf tubes and incubated with 500, 250 and 125 μM of different compounds, at 37 °C and 5% CO_2_ for 30 min. A non-treated control, 1% DMSO (vehicle control) and complete cell lysis control (Triton-X 2%) were included in each assay. After exposure samples were centrifuged at 4 °C at 1000 g for 15 minutes and 100 μl of supernatant transferred to a clean well in a 96-well, clear, flat-bottom plate (Corning Incorporated). Released haemoglobin was measured by absorption at 540 nm using Spectramax M5 plate reader. The experiments were performed in quadruplicate. The percentage of haemolysis was calculated as follows:

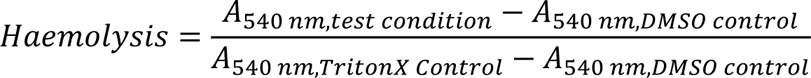

### Cell histology

HEK293 cells were plated in 24-well plates on exposure medium for 24 hours (500.000 cells/well). Cells were exposed to a subset of compounds at a lethal concentration of 20% (LC_20_). After 24-hours exposure, medium with compounds was removed and cells harvested by pipetting up and down in 250 μl of exposure medium. Cells from three wells treated with the same compound were pooled, and centrifuged for 5 minutes at 1.500 r.p.m. The cell pellet was fixed in 0.1 M cacodylate buffer containing 2% glutaraldehyde (v/v) (pH 7.4). After 1 hour on ice, the pellet was washed with 0.1 M cacodylate buffer and fixed in 0.1 M cacodylate buffer containing 1 % osmium tetroxide (w/v). After a further hour on ice the cell pellet was embedded in hard Epon and 1 μm slides were cut and stained with 1% toluidine blue and 1% borax (w/v). Histological abnormalities were assessed by microscopy using a Leica DM6B light microscope (Leica Microsystems B.V., Amsterdam, the Netherlands) at 100x magnification.

## Acknowledgements

Data collection experiments for the structures reported in the manuscript was carried at XALOC beamline at ALBA synchrotron (Cerdanyola del Valles, Spain) supported by ALBA BAG Proposal 2022075911. We acknowledge the ALBA synchrotron for provision of beam time and we would like to thank beamline staff for assistance. This project has received funding from the European Union’s Horizon 2020 research and innovation programme under the Marie Sklodowska-Curie grant agreement number 765147, and grants PID2019-108541GB-I00 from Spanish Government (Ministry of Science and Innovation) and PROMETEO/2020/012 by Valencian Government to A.M. We also thank the Utrecht Medical Centre (UMC) group from prof. Rob Willems for providing the enterococci.

## Supplemental material

**Supplementary Figure SF1.**
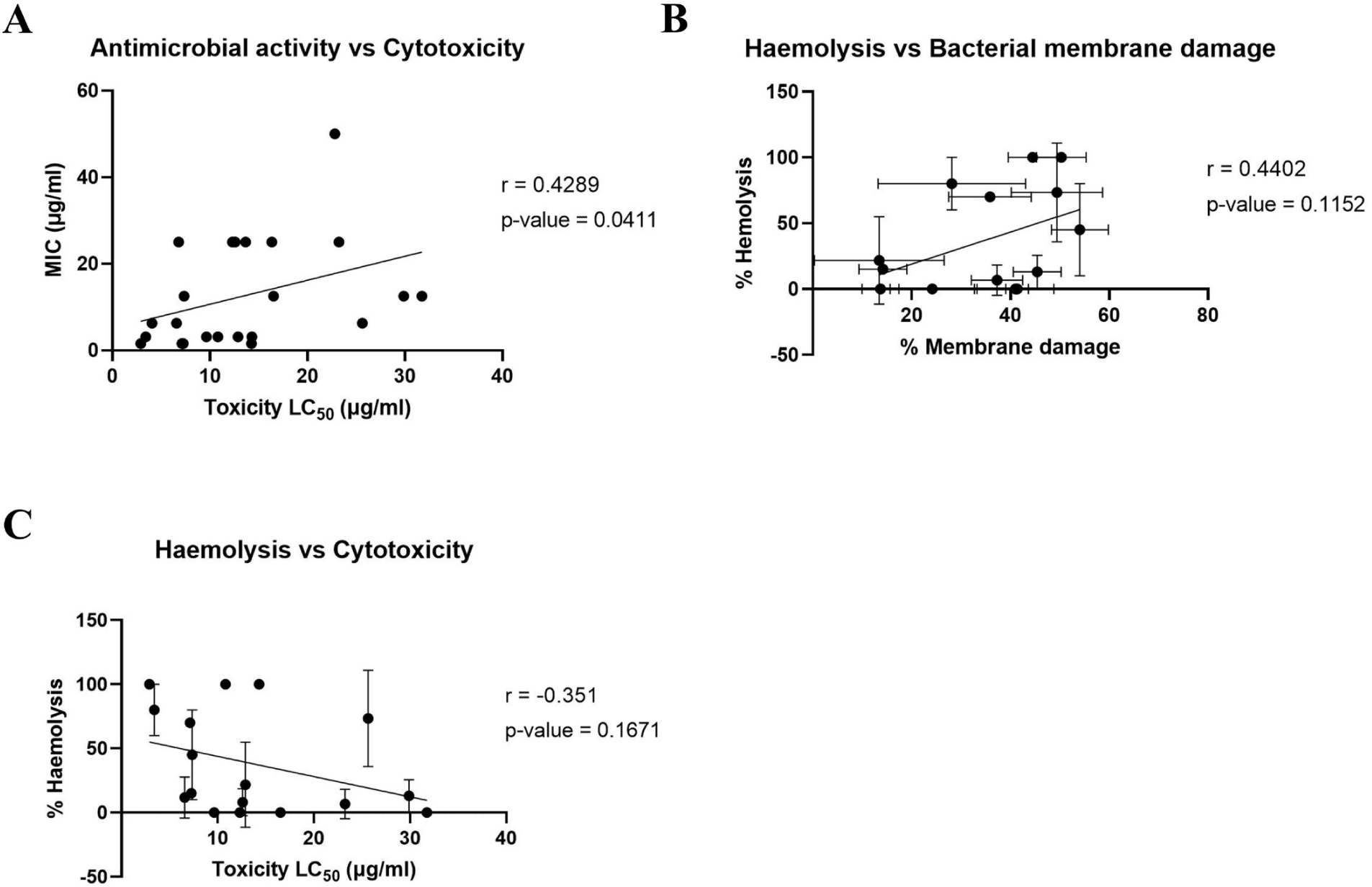
Correlation graphs of relevant parameters, including Spearman rank-correlation coefiecient (r) and p-value. **A.** Correlation between the antimicrobial activity expressed in MIC (μg/ml) and toxicity expressed as LC_50_ (μg/ml). **B.** Correlation between haemolysis percentage in sheep red blood cells and percentage of membrane damage (Suplementary Table 3). **C.** Correlation between percentage of haemolysis in sheep red blood cells and toxicity expressed as LC_50_ (μg/ml).

**Table ST1.** MIC data of DPP compounds against a panel of Gram-positive and Gram-negative strains, including *E. coli* outer membrane and efflux mutants. n.t. non-tested, *E. coli* JW5503 (Δ*tolC* mutant), *E. coli* D21f2 defective LPS core..

| DPP comp. | MIC ($\mu\text{g/ml}$) | | | | | | |
| --- | --- | --- | --- | --- | --- | --- | --- |
|  | *E. faecium* | *E. faecalis* | *P. haemolytica* | *P. aeruginosa* | *E. coli* | *E. coli* JW5503 | *E. coli* D21f2 |
| 1 | 12.5 | 12.5 | 25 | >250 | >250 | 1.56 | n.t. |
| 2 | 25 | 25 | 50 | >250 | >250 | 6.25 | 50 |
| 3 | >250 | >250 | 50 | >250 | >250 | 1.56 | n.t. |
| 4 | 3.12 | 3.12 | 125 | >250 | >250 | 6.25 | >50 |
| 5 | 12.5 | 12.5 | 12.5 | >250 | 50 | 3.12 | 50 |
| 6 | 6.25 | 6.25 | 12.5 | >250 | >250 | 3.12 | 25 |
| 7 | 3.12 | 3.12 | 25 | >250 | >250 | 3.12 | 25 |
| 8 | 3.12 | 1.56 | 250 | >250 | >250 | 12.5 | 25 |
| 9 | 25 | 25 | 25 | >250 | >250 | 12.5 | 50 |
| 10 | 250 | 250 | 62.5 | >250 | >250 | 6.25 | 250 |
| 11 | 12.5 | 12.5 | 62.5 | >250 | >250 | 1.56 | 25 |
| 12 | 50 | 50 | 62.5 | >250 | >250 | 12.5 | >50 |
| 13 | 25 | 25 | >250 | >250 | >250 | 12.5 | >50 |
| 14 | 12.5 | 12.5 | >250 | >250 | >250 | 3.13 | n.t. |
| 15 | 1.56 | 1.56 | >250 | >250 | >250 | 3.12 | n.t. |
| 16 | 50 | 50 | 50 | >250 | >250 | 3.13 | n.t. |
| 17 | >250 | 250> | 50 | >250 | >250 | 6.25 | n.t. |
| 18 | 50 | 50 | >250 | >250 | >250 | 6.25 | n.t. |
| 19 | 50 | 50 | >250 | >250 | >250 | 12.5 | n.t. |
| 20 | 25 | 25 | >250 | >250 | >250 | 6.25 | n.t. |
| 21 | 12.5 | 12.5 | 12.5 | >250 | >250 | 1.56 | >250 |
| 22 | 12.5 | 12.5 | 12.5 | >250 | >250 | 1.56 | 25 |
| 23 | 25 | 12.5 | 31.2 | >250 | 125 | 25 | 50 |
| 24 | >250 | >250 | 31.2 | >250 | 125 | 25 | 50 |

**Figure SF2.**
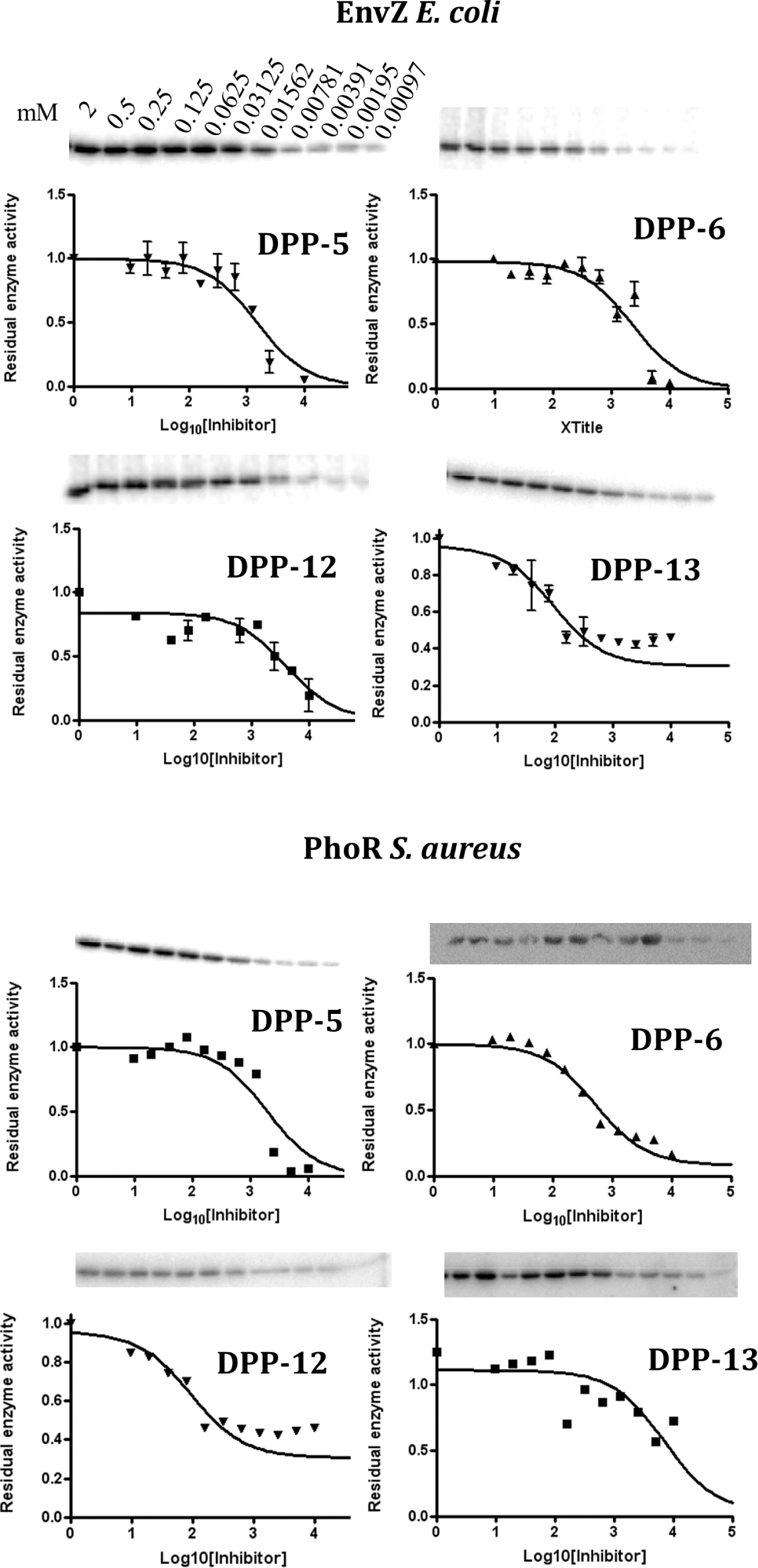
Autophosphorylation inhibition assays of DPP compounds against EnvZ form *E. coli* and PhoR from *S. aureus.* A range of concentrations of DPP compounds was tested for inhibition of autophosphorylation of two HKs (PhoR from *S. aureus* and EnvZ from *E. coli).* Amount of phosphorylated HK was detected via Western Blot and percentage of autophosphorylation calculated according to the positive (ATP) and inhibition (ATP+AMP-PNP) controls. Curves were plotted using Prism 4.1.

**Figure SF3.**
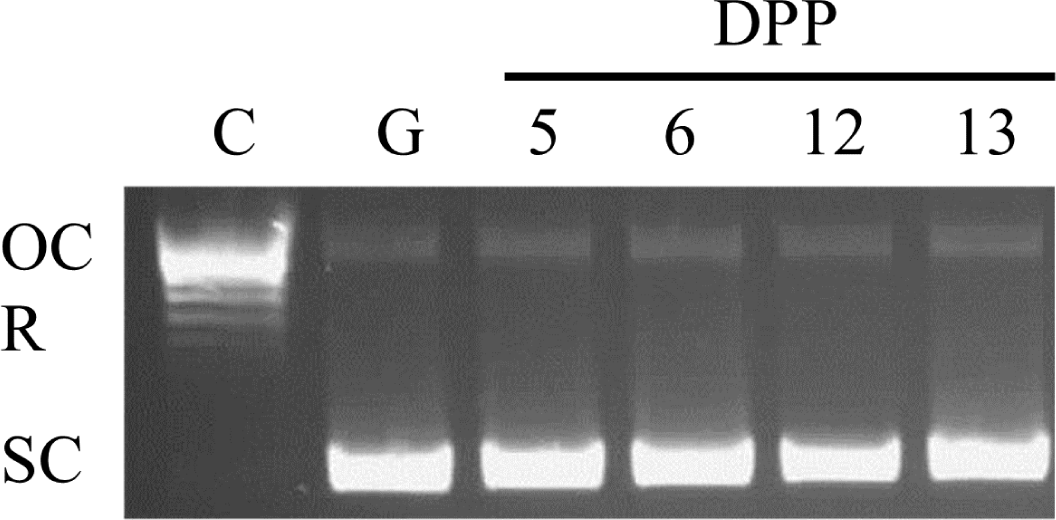
Bacterial gyrase inhibition assay of DPP compounds. Gyrase inhibition was tested using a supercoiling inhibition assay. Control (C) contained relaxed pBR322, Gyrase control (G) contains relaxed pBR322 and *E. coli* gyrase (0.1 mU/reaction). DPP-compounds were added at a concentration of 2 mM. No inhibition of the supercoiled activity of *E. coli* gyrase is observed by any of the DPP compounds. OC = nicked, open circular; R = relaxed topoisomers; SC = supercoiled topoisomers

## Supplementary method

### Gyrase inhibition assay

Gyrase activity inhibition assays were performed using *E. coli* gyrase supercoiling kit (Inspiralis, Norwich, UK)(57) following the manufacturer instructions. Briefly, assay buffer (35 mM Tris-HCl pH 7.5, 24 mM KCl, 4 mM MgCl_2_, 2 mM DTT, 1.8 mM spermidine, 1 mM ATP, 6.5% (w/v) glycerol, and 0.1 mg/ml albumin) was mixed with 0.5 μg relaxed pBR322, and 2 mM final concentration of desired inhibitor in DMSO. Reaction was started by adding 0.1 mU of *E. coli* gyrase per reaction, for a total final volume of 30 μl per reaction. Samples were incubated for 30 mins at 37 °C, stopped by adding 30 μl STEB (40% sucrose, 100 mM Tris-HCl pH 8.0, 10 mM EDTA and 0.5 mg/ml bromophenol blue) and 30 μl 24:1 chloroform/isoamyl alcohol and mixed by vortexing. Samples were centrifuged for 1 minute and 30 μl was loaded into a 1% agarose gel supplemented with SYBRSafe^TM^ followed by electrophoresis at 85 V for 2 hours. Gels were visualized using GelDoc XR+ gel documentation system (Bio-Rad).

**Suplementary Table ST2.** Porcentage of FITC-stained bacteria as measured using flow-citometry at different concentrations and different treatments.

| Treatment | % FITC stained<br>bacteria 0.5 xMIC | % FITC stained<br>bacteria 1 xMIC | % FITC stained<br>bacteria 2 xMIC |
| --- | --- | --- | --- |
| Unstained | 0 |  |  |
| Alive | 4.10 |  |  |
| Nisin | 16.37 | 53.32 | 59.41 |
| Novobiocin | 8.28 | 12.62 | 13.17 |
| 2 | 49.81 | 54.03 | 63.56 |
| 3 | 35.78 | 49.41 | 58.28 |
| 4 | 7.94 | 14.15 | 19.78 |
| 5 | 13.40 | 13.42 | 41.29 |
| 6 | 36.93 | 44.48 | 48.91 |
| 7 | 42.44 | 50.30 | 54.59 |
| 9 | 38.22 | 41.34 | 43.78 |
| 10 | 28.57 | 37.28 | 40.99 |
| 11 | 24.77 | 28.12 | 57.97 |
| 12 | 9.04 | 24.16 | 29.11 |
| 13 | 35.10 | 45.40 | 45.44 |
| 15 | 9.29 | 13.68 | 18.42 |
| 20 | 31.90 | 40.99 | 50.97 |
| 21 | 31.51 | 35.84 | 51.03 |

**Figure SF4.**
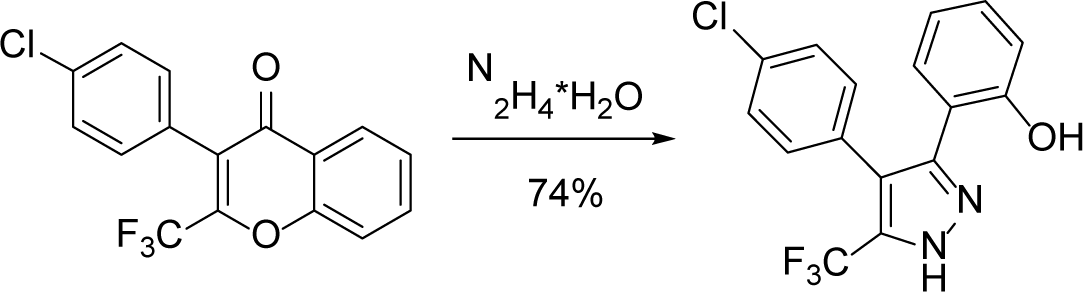
Schematic synthesis of DPP-22.

**Table ST3.**
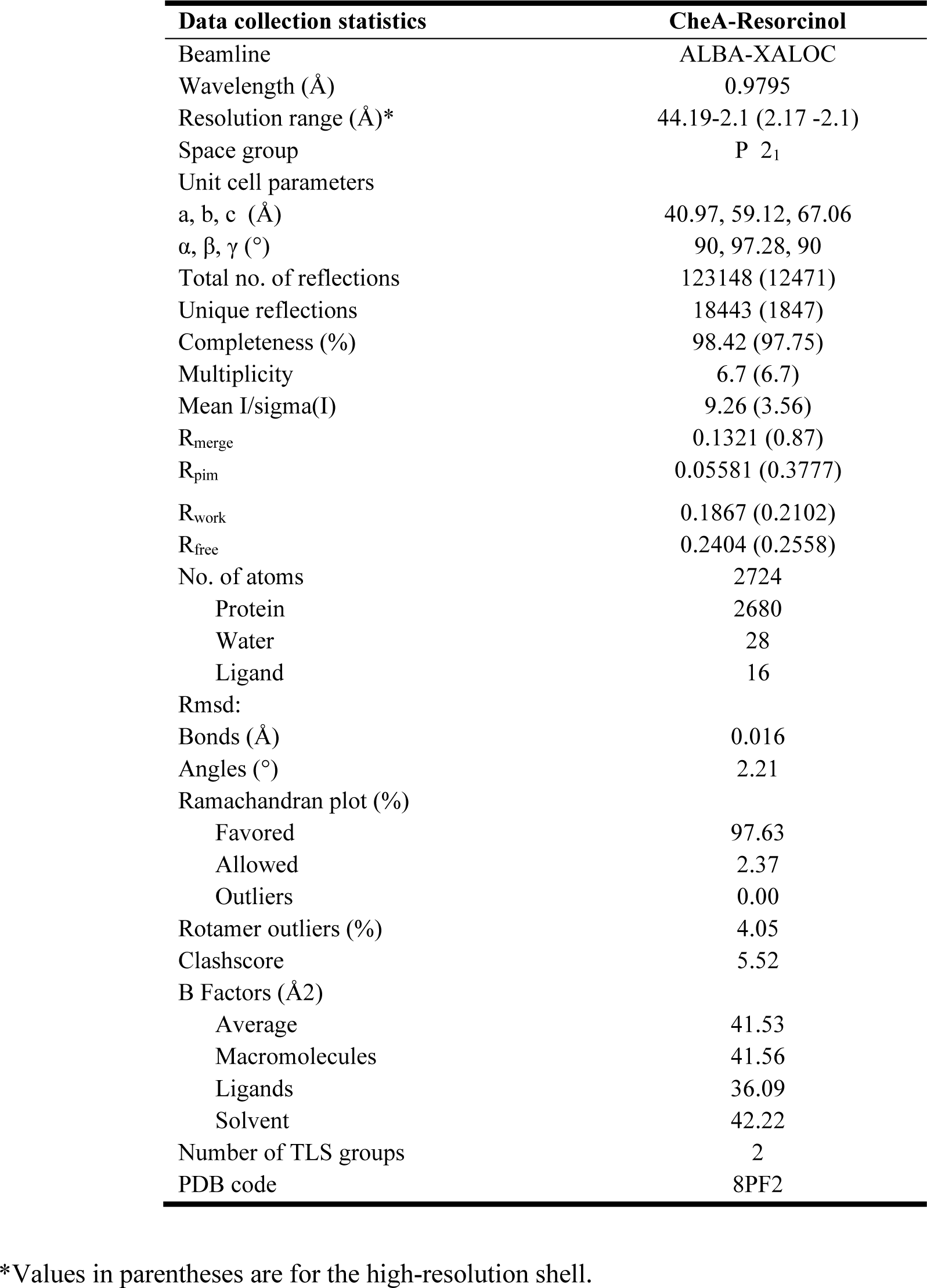
Data collection and refinement statistics for CheA-resorcinol complex.

| **Data collection statistics** | **CheA-Resorcinol** |
| --- | --- |
| Beamline | ALBA-XALOC |
| Wavelength (Å) | 0.9795 |
| Resolution range (Å)* | 44.19-2.1 (2.17 -2.1) |
| Space group | P 2<sub>1</sub> |
| Unit cell parameters |  |
| a, b, c (Å) | 40.97, 59.12, 67.06 |
| α, β, γ (°) | 90, 97.28, 90 |
| Total no. of reflections | 123148 (12471) |
| Unique reflections | 18443 (1847) |
| Completeness (%) | 98.42 (97.75) |
| Multiplicity | 6.7 (6.7) |
| Mean I/sigma(I) | 9.26 (3.56) |
| R<sub>merge</sub> | 0.1321 (0.87) |
| R<sub>pim</sub> | 0.05581 (0.3777) |
| R<sub>work</sub> | 0.1867 (0.2102) |
| R<sub>free</sub> | 0.2404 (0.2558) |
| No. of atoms | 2724 |
| Protein | 2680 |
| Water | 28 |
| Ligand | 16 |
| Rmsd: |  |
| Bonds (Å) | 0.016 |
| Angles (°) | 2.21 |
| Ramachandran plot (%) |  |
| Favored | 97.63 |
| Allowed | 2.37 |
| Outliers | 0.00 |
| Rotamer outliers (%) | 4.05 |
| Clashscore | 5.52 |
| B Factors (Å<sup>2</sup>) |  |
| Average | 41.53 |
| Macromolecules | 41.56 |
| Ligands | 36.09 |
| Solvent | 42.22 |
| Number of TLS groups | 2 |
| PDB code | 8PF2 |

\*Values in parentheses are for the high-resolution shell.

